# Production of antigenically stable enterovirus A71 virus-like particles in *Pichia pastoris* as a vaccine candidate

**DOI:** 10.1101/2023.01.30.526315

**Authors:** Natalie J Kingston, Joseph S Snowden, Agnieszka Martyna, Mona Shegdar, Keith Grehan, Alison Tedcastle, Elaine Pegg, Helen Fox, Andrew J Macadam, Javier Martin, James M Hogle, David J Rowlands, Nicola J Stonehouse

**Author notes:** Corresponding Authors: Nicola J. Stonehouse, and David J. Rowlands,.

## Abstract

Enterovirus A71 (EVA71) causes widespread disease in young children with occasional fatal consequences. In common with other picornaviruses, both empty capsids (ECs) and infectious virions are produced during the viral lifecycle. While initially antigenically indistinguishable from virions, ECs readily convert to an expanded conformation at moderate temperatures. In the closely related poliovirus, these conformational changes result in loss of antigenic sites required to elicit protective immune responses. Whether this is true for EVA71 remains to be determined and is the subject of this investigation.

We previously reported the selection of a thermally resistant EVA71 genogroup B2 population using successive rounds of heating and passage. The mutations found in the structural protein-coding region of the selected population conferred increased thermal stability to both virions and naturally produced ECs. Here, we introduced these mutations into a recombinant expression system to produce stabilised virus-like particles (VLPs) in *Pichia pastoris*.

The stabilised VLPs retain the native virion-like antigenic conformation as determined by reactivity with a specific antibody. Structural studies suggest multiple potential mechanisms of antigenic stabilisation, however, unlike poliovirus, both native and expanded EVA71 particles elicited antibodies able to directly neutralise virus *in vitro*. Therefore, the anti-EVA71 neutralising antibodies are elicited by sites which are not canonically associated with the native conformation, but whether antigenic sites specific to the native conformation provide additional protective responses *in vivo* remains unclear. VLPs are likely to provide cheaper and safer alternatives for vaccine production and these data show that VLP vaccines are comparable with inactivated virus vaccines at inducing neutralising antibodies.

## Introduction

Enterovirus A71 (EVA71) is one of the leading causes of hand, foot and mouth disease (HFMD), and is responsible for a considerable burden of disease in young children, especially in the Asia-Pacific region. While generally mild, severe cases can lead to neurological complications, including acute flaccid myelitis, meningitis, and death. Several vaccines based on inactivated virus have reached phase III clinical trial [1–5], with three different genogroup C4 vaccines being licenced for use in mainland China. These vaccines vary in total antigen content (Chinese Academy of Medical Sciences (CAMS), 100 AgU; Vigoo, 320 AgU; Sinovac, 400 AgU), each preventing more than 90% of EVA71 HFMD and all EVA71-associated neurological complications in stage 3 trials [6].

The impressive reduction in EVA71-associated illness in the 1- and 2-year follow up highlights the importance of vaccination, although there remain safety concerns surrounding the large-scale production of live virus prior to inactivation. Indeed, there have been several instances of inadvertent release of infectious poliovirus (PV) into the environment as a consequence of vaccine manufacture [7–12]. The implementation of alternative vaccine candidates, such as peptide, subunit or virus-like particle (VLP) vaccines has shown promising results in early-stage trials [13–20], and these approaches have improved safety as infectious material is not required during vaccine production. Indeed, VLPs produced in the yeasts *Pichia pastoris*, or *Saccharomyces cerevisiae* are routinely used in large-scale manufacturing of the licenced hepatitis B virus and human papillomavirus vaccines [21].

EV particles are assembled from 60 repeating copies of the viral structural proteins, VP0, VP3 and VP1. When these proteins assemble around the viral genome, the VP0 precursor is processed into VP4 and VP2. When initially produced during viral infection, both virions and empty capsids (ECs) are in the native (NAg) conformation. Several changes associated with genome encapsidation stabilise virions in the NAg conformation. These include VP0 cleavage and rearrangement of the internal protein network leading to increased intra- and inter-protomer interactions, increased inter-pentamer interactions, further enhancing particle stability [22–24]. The conversion from NAg to expanded state (HAg) is associated with several conformational changes including loss of a lipid molecule (pocket factor) from a pocket within VP1 at the base of the canyon [25]. At moderate temperatures in the extracellular environment, ECs rapidly convert to the HAg form, with a concurrent loss of pocket factor, changes within the internal protein network and a general expansion of the capsid. The current licenced EVA71 vaccines contain both virions and ECs, resulting in a combination of NAg and HAg particles.

For PV, only NAg particles are able to induce long-lived protective immunity and expanded HAg particles are ineffective as vaccines. In contrast, wild type (wt) EVA71 VLPs, which readily adopt the HAg conformation, induce protective responses in murine models [19]. Whether immunogenicity can be improved by using VLPs in the NAg conformation has yet to be established.

In PV, increased antigenic stability has been achieved through *in vitro* viral evolution studies, either by subjecting wt virus to successive rounds of heating and passage [26], or by growing a thermally sensitive mutant virus population at moderate temperatures [27]. In both instances, the passaged virus accumulated mutations within the structural proteins which provided increased resistance to heat. The introduction of combinations of these mutations into wt sequences resulted in thermally stable VLPs [28]. Using a similar approach, we have recently described the generation of a thermally stable genogroup B2 EVA71 virus population selected through successive rounds of heating and passage [29]. The selected mutations conferred resistance to antigenic conversion for both the virus and ECs [29]. Here, we have utilised this modified sequence to express recombinant stabilised VLPs (rsVLPs) in *P. pastoris*. We have characterised these rsVLPs antigenically and structurally and have compared immunogenicity to wt VLPs.

## Methods

### *Expression of EVA71 VLPs in* Pichia pastoris

EVA71 VLPs were produced in *P. pastoris* following previously described protocols with minor modifications [30]. Briefly, wt or rs P1 regions were amplified from a previously described reverse genetics systems [31]. The primers used for amplification encoded a 5’*Rsr*II restriction site and Kozak sequence (AAACGATG), the reverse primer introduced a stop codon and a 3’ *Fse*I restriction site. The wt and rs P1 regions were introduced into the pPink-HC vector under the AOX promoter and sequences were confirmed by Sanger sequencing. The PV 3CD region was shuttled into vectors at the *Sac*I restriction site, and plasmids introduced into and expressed in *Pichia* pink cells as described in Sherry *et al*. (2020) [30].

All solutions used in the production and purification of samples were generated with sterilised endotoxin free water, in addition sucrose solutions used for purification were supplemented with 0.02% sodium azide to prevent later endotoxin contamination. VLPs were isolated from *P. pastoris* cells by disruption at 40 kpsi and supplemented with 1 mM MgCl and 250 units denarase (c-LEcta) before incubation at room temperature for 2 hours with agitation. Samples were clarified at 4,000 rcf and clarified supernatant was precipitated overnight at 4°C with 8% w/v PEG-8000. Precipitated material was re-suspended in PBS, and insoluble material removed by centrifugation at 4,000 rcf followed by 10,000 rcf for 30 minutes. VLPs were pelleted through a 30% sucrose cushion at 150,000 rcf for 3.5 hours. Pellets were resuspended in 1 ml PBS and separated on a 15-45% sucrose gradient at 50,000 rcf for 12 hours. 1 ml fractions were collected manually (top down) and assessed for the presence of EVA71 proteins by western blot and ELISA [31].

For structural studies and immunisation trials VLPs underwent an additional cycle of gradient purification. Peak fractions (7-9) were diluted 1:2 to reduce the sucrose concentration, and underlaid with 25%, 30% and 45% sucrose steps. Samples were centrifuged at 150,000 rcf for 3 hours, fractionated and assessed for the presence of EVA71 reactive proteins by western blot and ELISA. Before immunisation samples were assessed for the presence of endotoxin using LAL test (Pierce). To remove sucrose, samples for structural studies underwent buffer exchange into PBS and were concentrated by centrifugation through a 100,000 MWCO PES concentrator as previously described [29].

### Western blot

Samples were prepared by mixing 1:1 (v/v) with 2x Laemmli buffer, denatured at 95°C and loaded on 12% (w/v) SDS-PAGE gels using standard protocols. Western blot analysis was carried out as previously described [29, 31]; separated samples were transferred onto PVDF and blocked (5% skimmed milk powder reconstituted in TBS with 0.1% Tween-20). EVA71 VP0/VP2 reactive proteins were detected using anti-EVA71 VP2 antibody clone 979 (Merck) at 1:2000 dilution and anti-mouse IgG HRP conjugate. Blots were developed using chemiluminescent substrate (Promega) and X-ray film.

### ELISAs

A sandwich ELISA method was utilised to determine the total and specific antigen composition of samples. To determine the total antigen content, we used mAb CT11F9 with 18/116 EVA71 antigen standard. A standard curve was generated from 2-fold diluted 18/116 standard and the resultant equation used to estimate the total antigen content of samples within a given plate [32]. To determine the specific NAg and HAg reactivity of samples we utilised a 16-2-2D scFv or mAb 979 in the detection phase as previously described [31]. Briefly, ELISA plates were coated overnight with polyclonal rabbit anti-EVA71 immune sera, plates were blocked, and samples were added to wells and incubated at 37°C for 1.5 hours. Either scFv or mAb was added to wells and incubated at 37°C for 1 hour. Anti-His HRP or anti-mouse HRP were used to detect scFv or mAb, respectively. Samples were incubated with OPD, reactions stopped with 3M HCl and the OD 492 nm measured using the Biotek PowerWave XS2 plate reader. Data was graphed using Graphpad Prism software.

### Electron microscopy

For negative stain EM, 3 μL of approximately 2000 AgU/ml VLP was applied for 30 s to carbon-coated 300-mesh copper grids (Agar Scientific, UK) following glow discharge of the grids (10 mA, 30 s). After wicking away excess liquid, grids were washed twice with 10 μL distilled H^2^O, then stained twice with 10 μL 2% uranyl acetate solution (UA). For the second application of UA, grids were incubated for 30 s, then lightly blotted and left to air dry. Imaging was performed using an FEI Tecnai F20 with field emission gun, operating at 200 kV and equipped with an FEI CETA camera, with a calibrated object sampling of 0.418 nm/pixel.

For cryoEM, 3 μL of the same sample of VLP was applied for 30 s to ultrathin continuous carbon-coated lacey carbon 400-mesh copper grids (Agar Scientific, UK) following glow discharge in air (10 mA, 30 s). Sample application was performed in a humidity-controlled chamber maintained at 8°C and 80% relative humidity. Excess liquid was removed by blotting using a range of blotting times (1.5 s to 3.5 s), then grids were vitrified in liquid nitrogen-cooled liquid ethane using a LEICA EM GP plunge freezing device (Leica Microsystems, Germany). Screening of grids and data collection was performed using an FEI Titan Krios transmission EM (ABSL, University of Leeds) operating at 300 kV. Imaging was performed with a calibrated object sampling of 0.91 Å/pixel (a complete set of data collection parameters are provided in Table S2).

### Image processing

Image processing was performed using Relion 3.1 [33, 34]. Motion-induced blurring was corrected using the Relion implementation of MotionCor2 [35], then CTFFIND-4.1 was used to estimate CTF parameters for each micrograph [36]. VLPs were picked with crYOLO using a model trained on a subset of manually picked particles [37]. Particles were extracted and 2× down-sampled for two rounds of 2D classification. Particles from selected 2D classes were re-extracted without down-sampling and subjected to 3D classification into two classes. The particles in each class were then processed separately, through 3D refinement using *de novo* initial models (with icosahedral symmetry imposed), further 3D classification without alignments, CTF refinement and Bayesian polishing. A final 3D refinement was performed with a mask to exclude solvent and calculation of solvent-flattened Fourier shell correlations (FSCs). Sharpening was performed using a solvent-excluding mask, and the nominal resolution was calculated using the ‘gold standard’ FSC criterion (FSC = 0.143) [38]. Local resolution was determined using Relion.

### Model building and refinement

To build an atomic model for the VLP the PDB-3VBS model was used and was rigid-body fitted into a single asymmetric unit within the density map using UCSF Chimera [39]. The model was altered to represent the rsVLP sequence in coot (sequence described in Kingston e*t al*., 2022 [29]) and regions of the peptide backbone without supporting density were removed. Iterative cycles of inspection and manual refinement in Coot [40], followed by automated refinement in Phenix [41], were performed to improve atomic geometry and the fit of the model within the density. To avoid erroneous positioning of the model within density corresponding to adjacent protomers, density from adjacent asymmetric units was occupied with symmetry mates during automated refinement. Molprobity was used for model validation [42].

### Structure analysis and visualisation

Visualisation of structural data was performed in UCSF Chimera [39], UCSF ChimeraX [43] and PyMOL (The PyMOL Molecular Graphics System, Version 2.1, Schrödinger, LLC).

### Cells and viruses

An EVA71 reverse genetics system of strain MS/7423/87 was generated as previously described [31]. HeLa cells were obtained from the National Institute of Biological Standards and Control (NIBSC), Vero cells were obtained from ATCC and human rhabdomyosarcoma (RD) cells were obtained from CDC by NIBSC.

### Virus preparation for immunisation

Four confluent 75cm^2^ flasks of RD cells were infected, cell sheets were then frozen at −20° and thawed, and cell debris removed by centrifugation. Igepal was added to supernatants to a final concentration of 0.1% and viral particles were concentrated by centrifugation at 4°C through a 10ml 30% sucrose cushion made up in 50mM NaCl and 10 mM Tris HCl pH7.2. Pellets were resuspended in 0.5ml 6-salt PBS, layered on 10ml 15%- 30% sucrose gradients in lysis buffer (6-salt PBS containing 0.5% sodium deoxycholate, 20mM EDTA and 1% Igepal) and centrifuged at 4°C for 12h at 18,000rpm in a Beckman SW41 rotor before harvesting into 18 0.5 ml fractions. Fractions were screened by ELISA to identify virus and EC peaks.

### Immunisation

10 groups of 8 female BALB/c mice were immunised intraperitoneally twice, 3-weeks apart with 100 AgU EVA71 antigen or PBS in the presence or absence of Alhydrogel (0.2%). Serum samples were collected via the tail vein on days −1 and 20, terminal bleeds were collected by cardiac puncture while mice were under terminal anaesthesia 14 days after the final vaccine dose was administered.

### Ethical approval

Animal experiments were performed under license PPL P4F343A03 granted by the UK Home Office under the Animal (Scientific Procedures) Act 1986 revised 2013 and reviewed by the internal NIBSC Animal Welfare and Ethics Review Board. Animals were housed under specific pathogen-free conditions.

### Neutralisation assay

Virus neutralisation assays were carried out according to the Pharmacopeial method established at NIBSC previously [44]. Briefly, 2-fold serial dilutions of antisera were generated and combined with 100 TCID_50_ of type-specific EVA71 (genogroup B2) and incubated at 35°C for 3 hours before the addition of 1×10^4^ RD cells. Plates were incubated at 37°C for 6 days, wells were fixed and stained with naphthalene black solution in acetic acid. Endpoint neutralisation titres were calculated using the Spearman-Karber method.

### Specific anti-EVA71 titres

The total reactive antibody contents of serum samples were assessed against genogroup B2 virions and VLPs. Briefly, ELISA plates were coated and blocked as above, 100 AgU (50 μl or 200 AgU/ml) of virions, wtVLPs or rsVLPs were added to wells and incubated at 37°C for 1 hour. Plates were washed and serum dilutions were added to wells in duplicate in a final volume of 50 μl. After a further 1 hour incubation at 37°C plates were washed and 50 μl 1:2000 diluted cross-adsorbed anti-mouse HRP (Sigma) was added to each well and incubated for 1 hour at 37°C. Plates were again washed before the addition of 100 μl Sigma-fast OPD (Sigma) for 15 minutes in the dark at ambient temperature. Reactions were stopped with the addition of 50 μl 3M HCl and the optical density recorded at 492 nm.

## Results

### *Generation of EVA71 VLPs in* Pichia pastoris

EVA71 NAg and HAg VLPs were produced using the *P. pastoris* expression system, as described previously. Heat stressing of samples of EVA71 virus resulted in a population in which both virions and ECs had enhanced thermal stability [29]. The mutations responsible for stabilisation were introduced into the P1 structural protein-coding region and both this modified sequence and the wt sequence were cloned into dual promoter plasmids to co-express the P1 region and PV 3CD (to facilitate processing of the P1 precursor protein) (Fig 1A). The plasmids were electroporated into *Pichia* pink for VLP expression (termed recombinant stabilised, rsVLPs). Cell lysates were purified on 15-45% sucrose gradients and fractions assessed by western blotting for the presence of VP0, detected using mAb 979 (Fig 1B).

**Figure 1:**
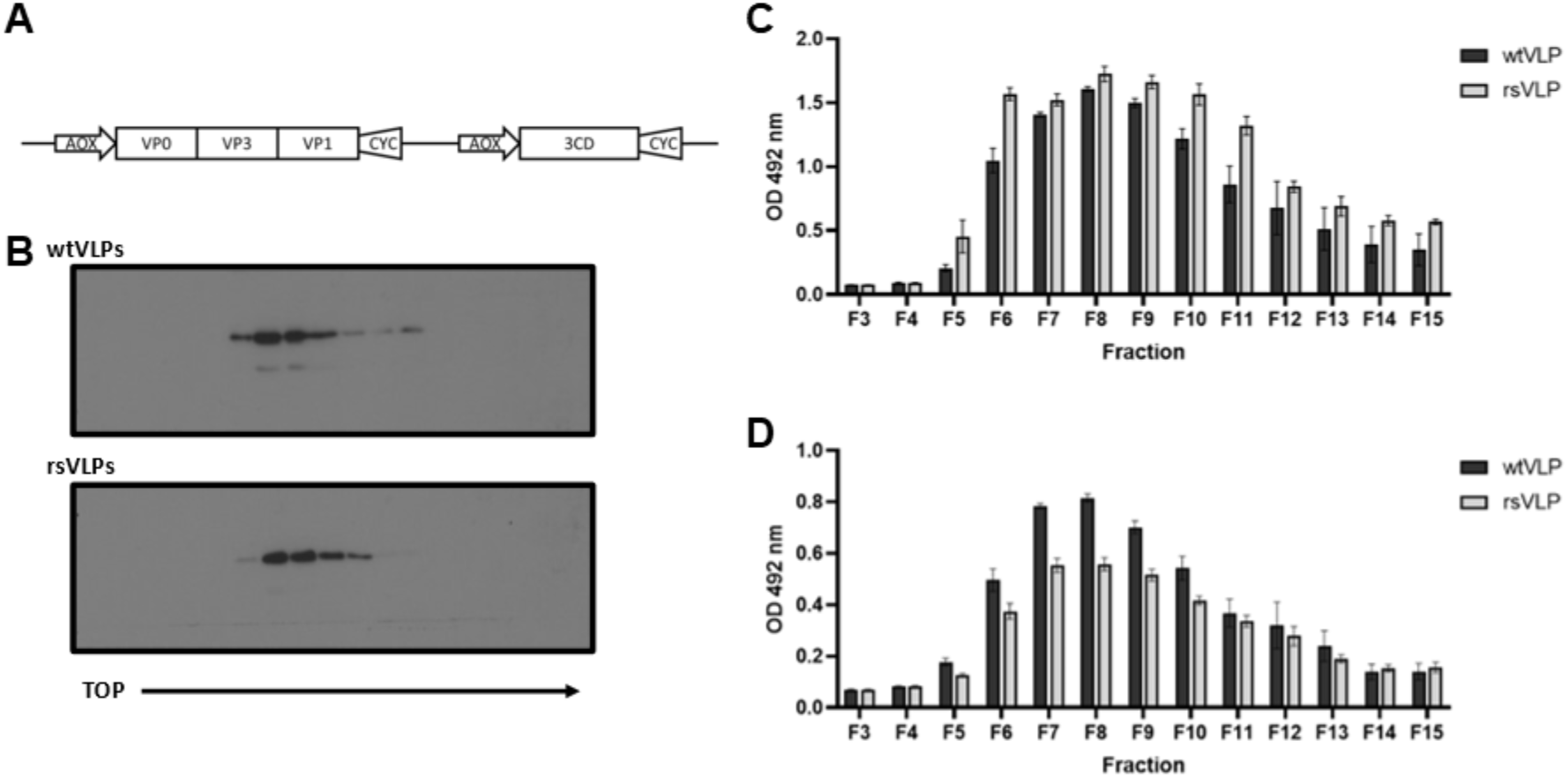
Expression of EVA71 VLPs. (**A**) EVA71 VLPs were produced in *P. pastoris* using the dual promoter expression cassette depicted. wt and rs VLPs were purified on 15-45% sucrose gradients. (**B**) Fractions were assessed by western blot using mAb 979, a representative image of n = 3 is shown. (**C**) Fractions were diluted 1:10 and heated to 55°C to facilitate conversion to a HAg conformation. Samples were assessed for reactivity with CT11F9, graph shows mean raw OD 492 nm ± SEM, n = 3 in duplicate. (**D**) Unheated fractions were diluted 1:10 and assessed for reactivity with HAg-specific mAb 979, graph shows mean raw OD 492 nm ± SEM, n = 3 in duplicate.

The total antigenic content of VLP gradient fractions was assessed by ELISA using the monoclonal antibody CT11F9. This antibody recognises both NAg and HAg conformations of EVA71 and is used as an international standard for EVA71 antigen quantification. We previously showed that both wt and rs ECs completely convert to the HAg conformation at 55°C [29] and the VLP sucrose gradient samples were heated to this temperature to negate the potential difference in CT11F9 reactivity with NAg and HAg particles. Following this treatment, CT11F9 reacted well with both wtVLPs and rsVLPs (Fig 1C). From 200 ml of yeast culture, the total antigen content within the peak fractions (7-9) was determined by comparison to the 18/116 antigen standard [32]. Both wt and rsVLPs contained similar levels of total antigen (Fig S1), with approximately 70 (CAMS 100 AgU/dose) human vaccine doses in the peak fractions for each VLP type [6].

In addition, we used mAb 979 in the detection phase of ELISA to determine whether the particles present in the gradient fractions had equivalent HAg content. Although both particle types reacted well with mAb 979 (Fig 1D), wtVLPs had greater reactivity than rsVLPs indicating a difference in antigenic composition.

### Antigenic stability of VLPs

To understand the antigenic composition of wtVLPs and rsVLPs more precisely, we performed antigen conversion assays which can detect conformational change within particles. The loss of NAg reactivity should be associated with a coincident gain in HAg reactivity, and these changes can be induced through receptor engagement, as well as by alterations in pH, ions, and temperature. We elected to use heat to induce particle expansion, which was assessed using ELISAs specific for HAg (mAb979, Fig 2A) or NAg (16-2-2D scFv, Fig 2B) to measure change in antigen composition [31]. The reactivity of wtVLPs with the HAg-specific antibody mAb 979 did not change significantly on heating. However, a greater than two-fold increase in signal was seen when the temperature of the rsVLPs sample was raised from 4°C to 50°C (Fig 2A). There was a small reduction in reactivity of wtVLPs with the NAg-specific 16-2-2D scFv after heating, suggesting the presence of some NAg in the wtVLPs (Fig 2B). The NAg reactivity of the rsVLPs was significantly more resistant to heating compared to wtVLPs, with more than 90% of signal retained at 40°C, although higher temperatures resulted in greater loss of NAg reactivity (Fig 2B), and concurrent gain in HAg reactivity (Fig 2A).

**Figure 2:**
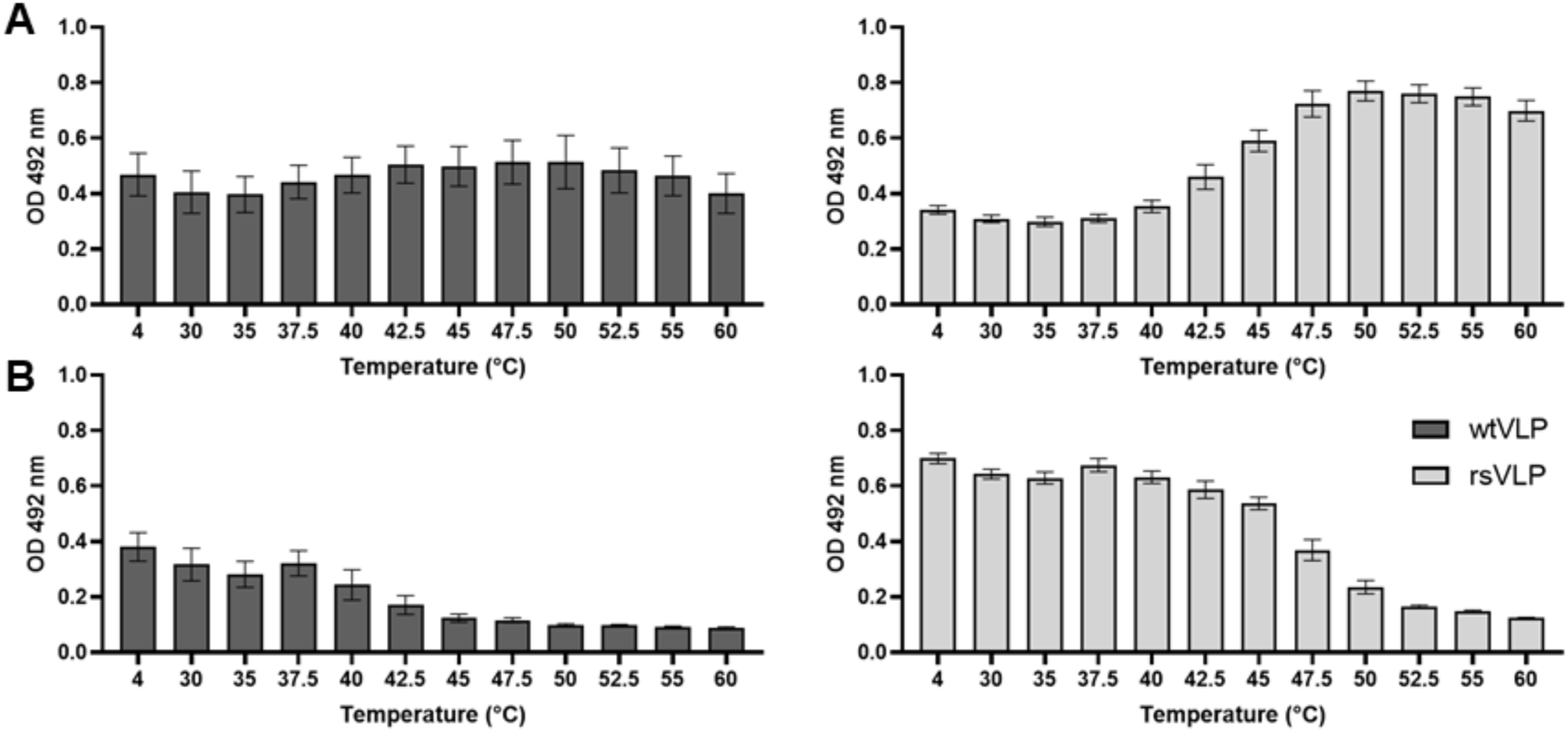
Antigenicity of EVA71 VLPs. Characterisation of the thermal stability of VLPs produced in *P. pastoris*. VLPs were incubated at a range of temperatures (x-axis) and assessed for the presence of (**A**) HAg reactivity using mAb 979 and (**B**) NAg reactivity using the 16-2-2D scFv. Graphs show mean raw OD 492 nm ± SEM, n = 3 in duplicate.

To more precisely determine the proportion of rsVLP in the HAg conformation (and indirectly determine the proportion in the NAg conformation) we made the following assumptions: 1) the sample at 4°C contains the maximum amount of NAg, 2) all particles have converted to HAg after incubation at 55°C and 3) any observed gain in HAg reactivity is as consequence of particles converting from NAg to HAg.

A standard curve can be generated to determine the loss of signal associated with a *n*-fold reduction in total particle number (Fig S2). By comparing the HAg-reactive signal from particles incubated at 4°C and 55°C, it is possible to calculate the proportion change in HAg (Fig S2). Accordingly, we identified the proportion of NAg and HAg particles present within the VLP samples at each of the temperatures used (Fig 3 & Table S1). By using the proportion of NAg and imposing a linear regression within the dynamic range (40-50°C) we can estimate the temperature at which 25%, 50% and 75% of NAg particles are converted to the HAg form (CT_##_) (Fig 3 & Fig S3). Together these data indicate that approximately 87% of rsVLPs produced in *P. pastoris* are in the NAg conformation and have a CT_50_ of approximately 44°C (Fig 3).

**Figure 3:**
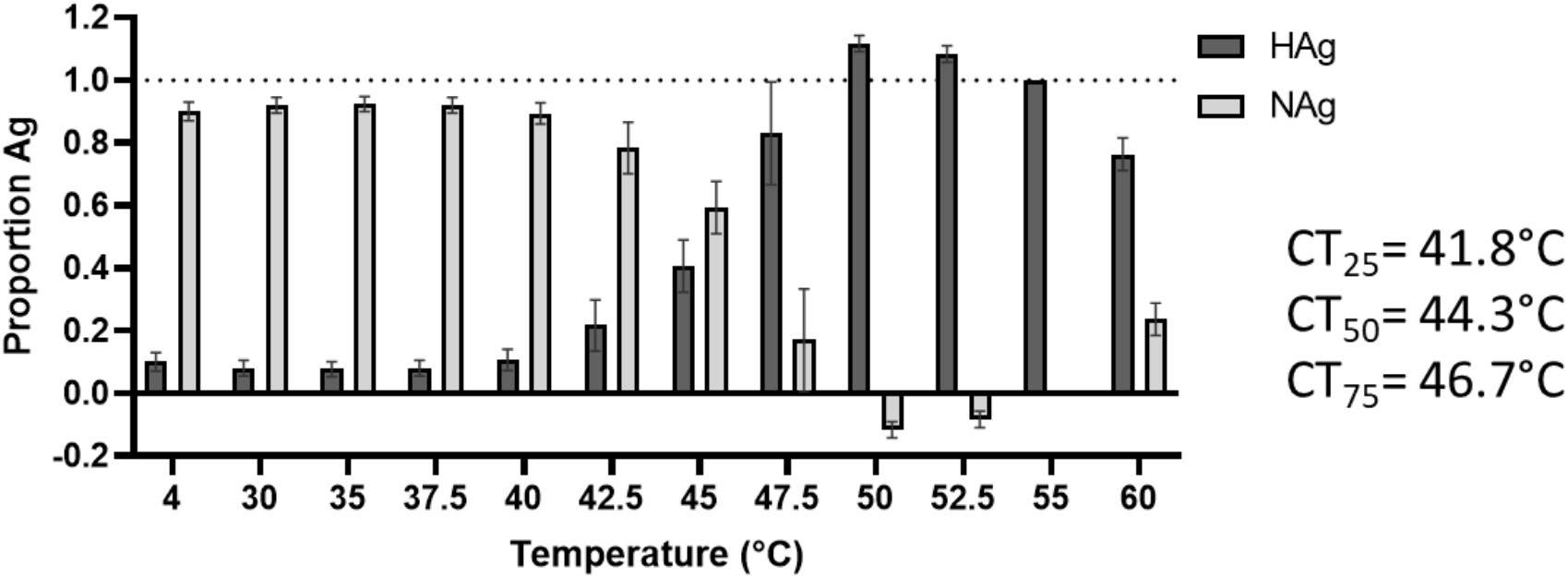
Antigenic stability of rsVLPs. The proportion of HAg present within the rsVLP samples was determined using the equation *proportion HAg* = *n^(OD 4°C-OD 55°C)/incline^*. The NAg proportion was calculated as the inverse of this number and the conversion from NAg to HAg graphed across a range of temperatures (x-axis). The temperature at which 25%, 50% and 75% of NAg is converted to HAg is indicated (CT_##_). Graph shows mean ± SEM, n = 3.

### Structure of rsVLPs

A number of mutations potentially contribute to stabilisation of the rsVLPs in the NAg conformation, including VP3-I235M, VP1-Y116C, VP1-K162I and VP1-P246A (described in Kingston *et al*. (2022) [29]). However, the mechanism by which these mutations contribute to antigenic stabilisation was unclear. To understand the structural consequences of these mutations and gain insight into the mechanism of stabilisation, we determined the structure of rsVLP to high resolution using cryoEM.

VLPs were generated as described above and subjected to a subsequent sucrose gradient for additional purification. To remove sucrose from samples, peak fractions were concentrated across a 100kDa MWCO spin concentrator and diluted into PBS, before being assessed by negative stain TEM. Both wt and rsVLPs showed morphologies consistent with EV particles (Fig S4). rsVLPs were also assessed by cryoEM.

In anticipation that a subset of the rsVLPs would have converted to an HAg conformation, cryoEM data was subjected to 3D classification into two classes. Based on visual inspection of the resultant density maps the major class, which was resolved to 2.4 Å, contained ~90% of particles and was clearly consistent with the NAg form of EVA71. However, the minor class (~10% of particles), which was resolved to 3.4 Å, was more challenging to classify as structurally native because several features were reminiscent of both NAg and HAg particles. This minor class showed weak density in several regions, including that corresponding to the N-terminal region of VP0 (residues corresponding to VP4), fewer resolved residues at the VP1 N-termini and reduced resolution around the icosahedral two-fold axis, all indicative of an HAg form (Fig 4). However, both populations contained particles with a similar diameter (approximately 30.6 nm) and had density maps which correlated by more than 95% when displayed at a 1σ threshold.

**Figure 4:**
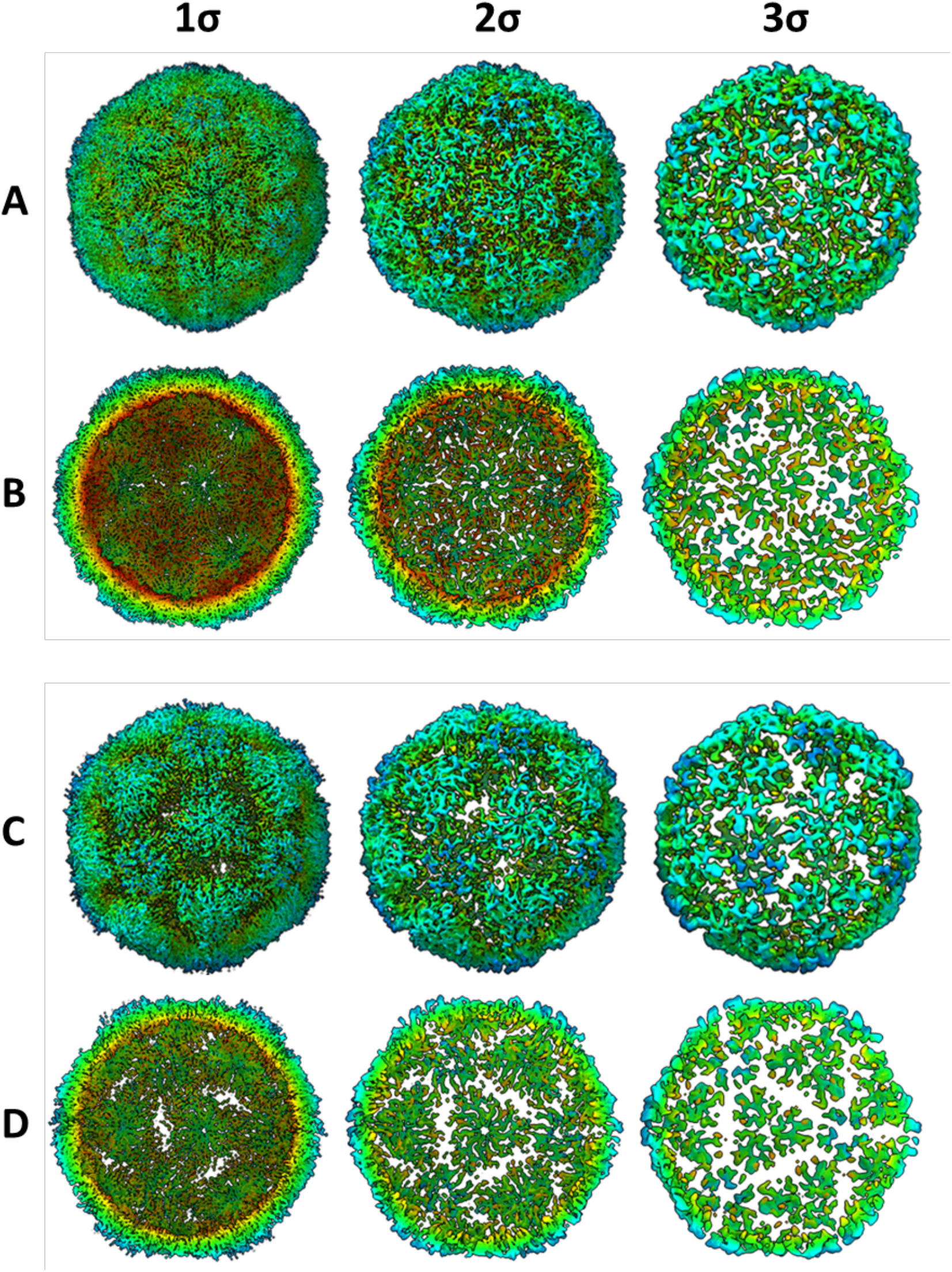
Dual conformation of rsVLPs. Complete (**A** & **C**) and sectional isosurface representation, with the front half of the particle removed to show the inner surface of the capsid shell (**B** & **D**) of 2.4 Å (major population; A and B) and 3.4 Å (minor population; **C** & **D**) rsVLP map, contoured at different thresholds (1 σ–3 σ).

Intriguingly, where resolution was sufficient to permit analysis, the minor class shared many of the structural features associated with the NAg conformation, including the presence of density in the VP1 pocket, consistent with the presence of a pocket factor (Fig S5). Consequently, it is not clear whether the minor population represented an alternative structural form of EVA71, an intermediate conformation in the transition from NAg to HAg, VLPs with defects, or partially disassembled particles.

Unlike high-resolution mature virion structures of EVA71 (PDB-3VBS), the rsVLPs in the NAg conformation lack resolution for residues 1-57 of VP1. The presence of resolved residues 57-72 along the internal network of the particle within the major rsVLP reconstruction may contribute to stabilisation of the inter-pentamer interface. These stabilising interactions are not seen within canonically HAg particles (PDB-3VBU), or the minor population rsVLPs resolved here, although it is unclear if the latter is a consequence of conformational change or lower local resolution.

The mutated residues present in the rsVLPs are located along the upper surface of the canyon, and in several regions, which show large conformational differences between the canonical NAg and HAg models (Fig 5). It is likely that the mutations function cooperatively to provide antigenic stability in the NAg form to the rsVLPs.

**Figure 5:**
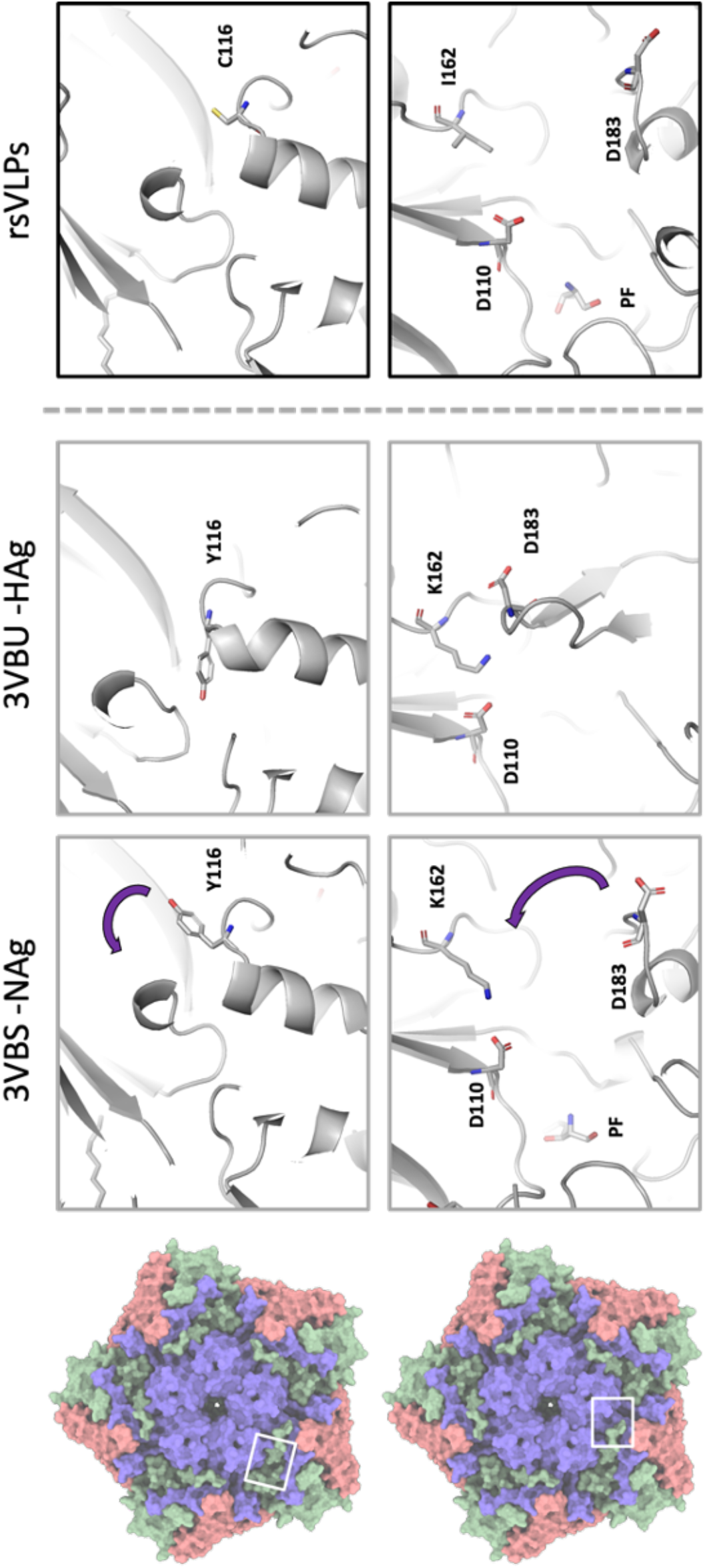
Structural characterisation of rsVLPs. Depiction of different conformations observed with NAg to HAg shift, and the loss of pocket factor. Cartoon and stick representation of the native conformation (PDB-3VBS), expanded conformation (PDB-3VBU) and rsVLP (major population) focusing upon the region surrounding the canyon and pocket. Residues associated with EC VLP stabilisation in rsVLPs are highlighted: VP3 I325M, VP1 Y116C, VP1 K162I, and the interacting residues VP1 D110, and VP3 D183. Some regions of the capsid have been clipped for clarity.

VP1 Y116 is localised at the upper surface of the canyon between the VP1 B- and C-strands. In the wt virus, rotation of the aromatic sidechain is associated with the separation of these stands, β-barrel expansion and loss of the pocket factor. It is possible that the Y116C mutation makes this expansion less energetically favourable (Fig 5). The precise role of VP1 K162I is less clear and may act both to prevent loss of pocket factor (Fig 6) and also to reduce the stability of the HAg form through its inability to make a salt bridge with D183 in the VP3 GH-loop (Fig 5).

**Figure 6:**
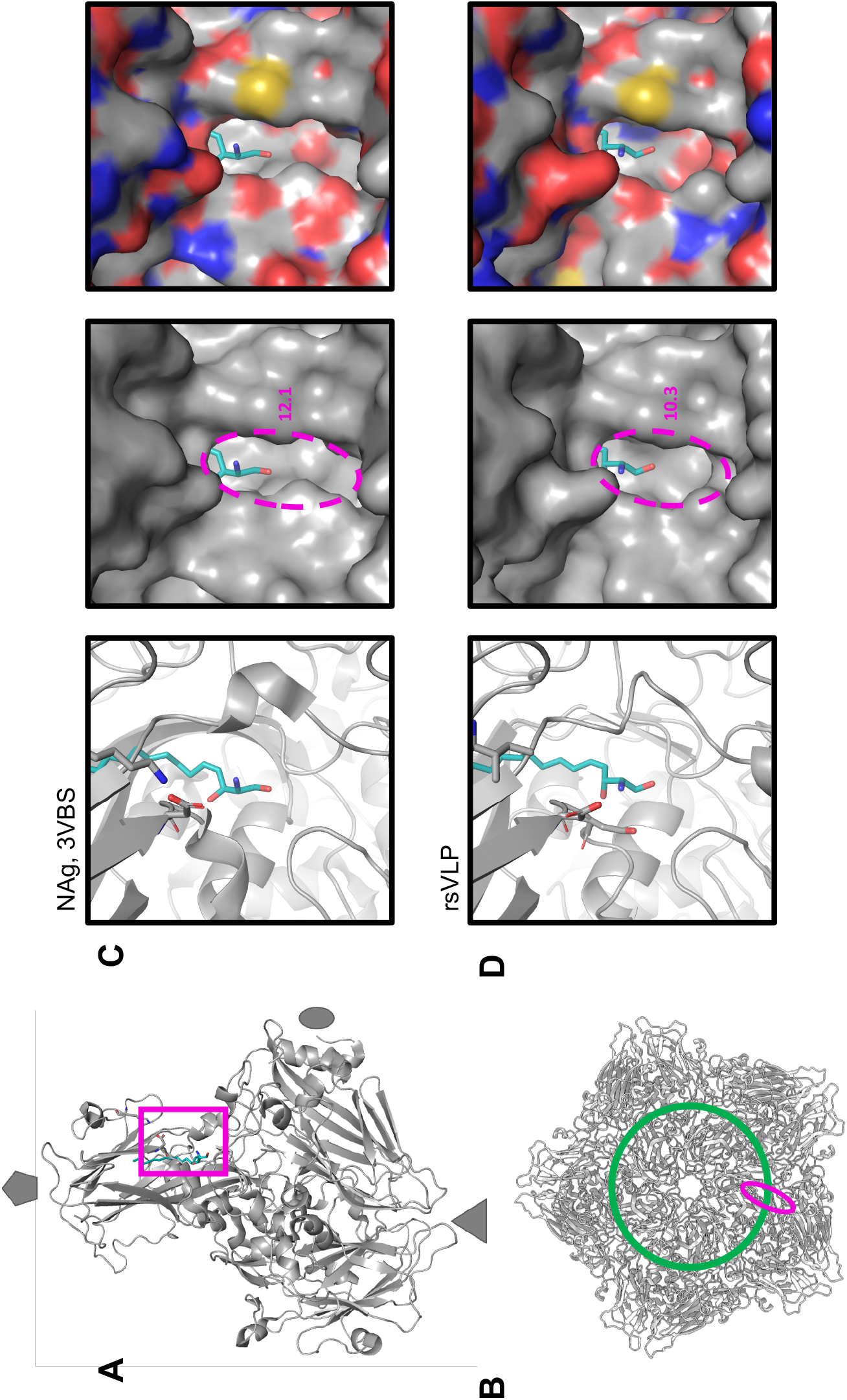
Narrowing of pocket opening in rsVLPs. Cartoon and surface representation of the capsid region surrounding the canyon and pocket. **A**) The approximate position of the displayed region in the context of an asymmetric unit is shown with a 50° rotation along the y-axis relative to the position shown in the panels on the right. **B**) The approximate position of the mouth of the pocket depicted as a pink ellipse relative to the floor of the canyon (green line) in the context of a protomer. Cartoon and surface representation of the capsid region surrounding the canyon and pocket within **C**) NAg virus and **D**) rsVLPs. Cartoon representation displaying pocket factor (teal), VP1 D110 and VP1 K162 or I162 as sticks. For clarity, the local surface is represented in both greyscale (middle) and coloured with respect to local atoms (right), oxygen; red, nitrogen; blue, sulphur; yellow.

A series of minor conformational changes around the pocket appear to result in a substantial reduction in the diameter of the pocket mouth (Fig 6). The absence of the salt bridge between VP1 K162 and D110 changes the position of the carboxylate groups of D110, resulting in several subtle changes within this region. A minor narrowing of the mouth running parallel with the canyon floor (depicted as a green circle in Fig 6) is observed in the rsVLPs compared to the canonical NAg particles (PDB-3VBS). The opening of the mouth of the pocket running perpendicular to the canyon shows a far more drastic narrowing, where the D110 carboxylate group protrudes an additional 1.1 Å over the canyon (Fig 6). The opening at the mouth of the canyon can be approximated as an ellipse, and these changes reduce the diameter of this along the major axis from 12.1 Å in the PDB-3VBS structure to just 10.3 Å in the rsVLPs, amounting to approximately 15% reduction in the space available for pocket factor egress, likely contributing to the increased antigenic stability of the rsVLPs.

### Immunisation

To compare the immunological responses to NAg or HAg conformations of EVA71 particles, mice were immunised twice with 100 AgU 18/116 inactivated genogroup C4 control virus (18/116), genogroup B2 virus, wtVLP, rsVLPs, or PBS in the presence or absence of Alhydrogel. Sera were assessed for the presence of antibodies specific for wtVLPs and rsVLP (Fig 7), for antibodies against genotype B2 virus (Fig 8A), and for neutralising antibodies against genotype B2 virus (Fig 8B).

**Figure 7:**
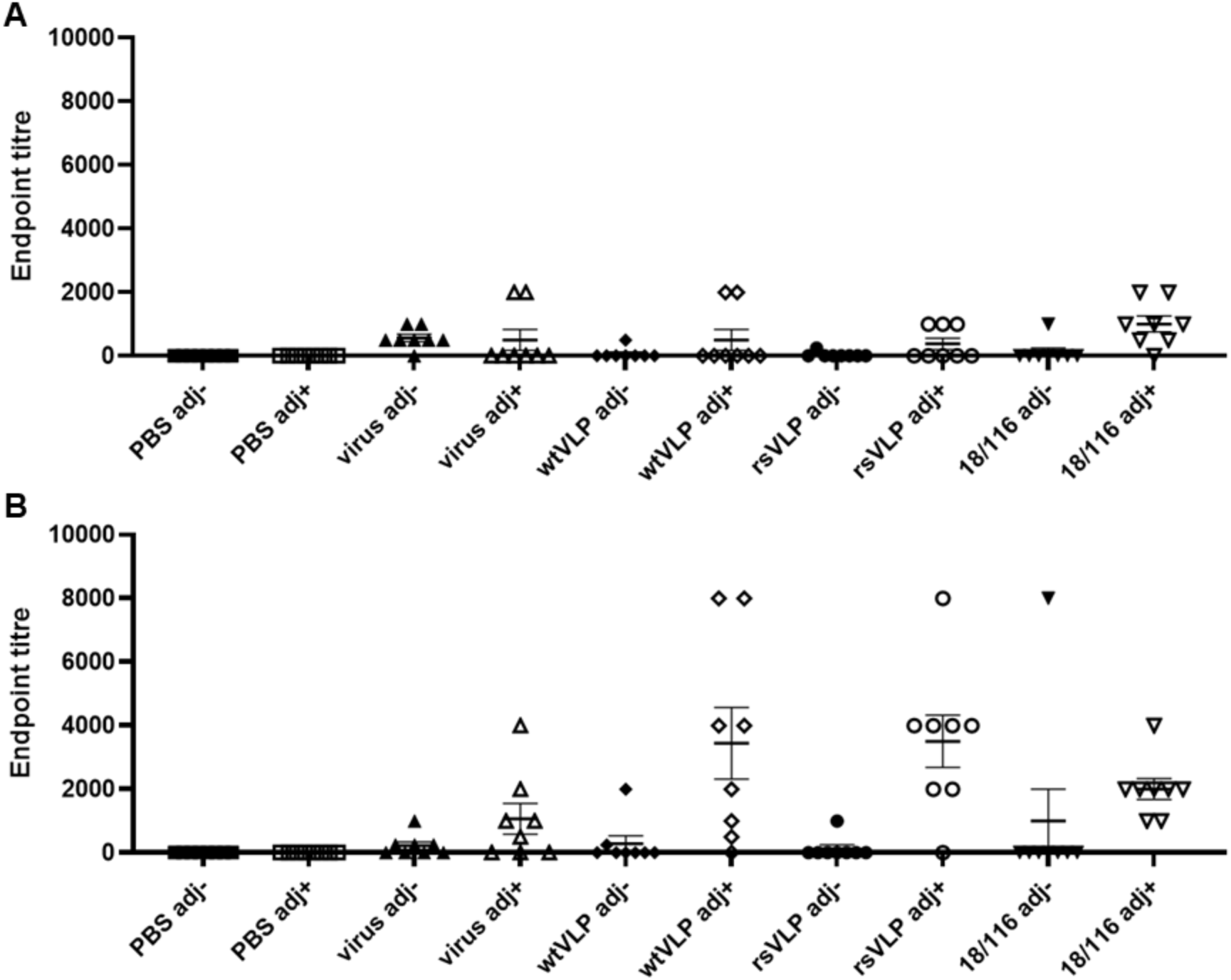
Reactivity with wt and rsVLPs by ELISA. Sera were collected from mice immunised twice with PBS or 100 AgU of EVA71 particles in the presence or absence of adjuvant (adj), indicated along the x-axis. Serum samples were then tested for reactivity with (**A**) wtVLPs or (**B**) rsVLPs. To each well, approximately 200 AgU of VLP was added, and sera were assessed at dilutions between 1:250 and 1:8000 in duplicate. Samples were considered positive if the average OD was >2x that of the negative (no serum) control wells within each plate. Graphs show the endpoint titre for individual animals, mean ± SEM, n = 8 in duplicate (n = 7 for PBS adj+ group). Abbreviations: adj-; unadjuvanted immunogen, adj+; adjuvanted immunogen, virus; genotype B2 virus immunogen, 18/116; genogroup C4 virus immunogen.

**Figure 8:**
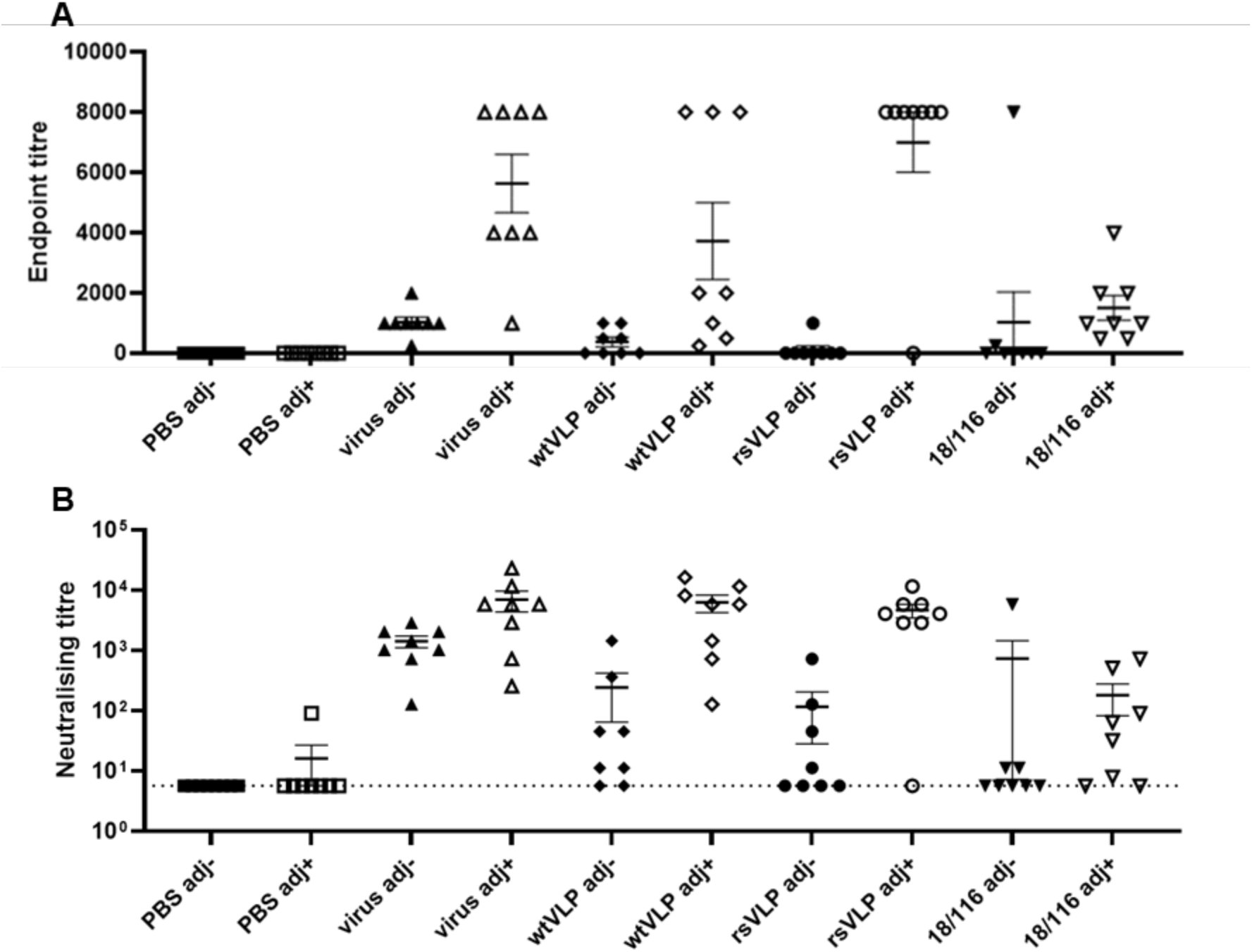
Total reactive and neutralising titres against virus. In addition, the sera were also used to assess reactivity with virus samples (identical serum samples as used above). (**A**) Serum samples were tested for reactivity with virus by ELISA. To each well, approximately 200 AgU of virus was added, and sera were assessed at dilutions between 1:250 and 1:8000 in duplicate. Samples were considered positive if the average OD was >2x that of the negative (no serum) control wells within each plate. Graph shows the endpoint titre for individual animals, mean ± SEM, n = 8 in duplicate (n = 7 for PBS adj+ group). (**B**) Virus neutralisation was assessed against 100 TCID_50_ of genogroup B2 EVA71, neutralising titres were determined using the Spearman-Karber method. Graph shows mean titres from individual animals, as well as group mean ± SEM, n = 8. Abbreviations: adj-; unadjuvanted immunogen, adj+; adjuvanted immunogen, virus; genotype B2 virus immunogen, 18/116; genogroup C4 virus immunogen.

Antibody responses that detect wtVLPs were relatively low in each of the immunisation groups, in contrast to the responses against rsVLPs (Fig 7). The rsVLPs performed better despite similar quantities of VLPs being captured in these ELISAs, as determined by reactivity with CT11F9 (Fig S6). Antibody titres induced by immunisation with wtVLPs or rsVLPs and detecting using rsVLPs were higher in the presence of adjuvant compared to the unadjuvanted groups (p = 0.0159 & p = 0.0012, respectively), although this adjuvant-specific improvement in immunogenicity was less pronounced for the B2 virus and 18/116 immunisation groups (p = 0.1139 & p = 0.3580, respectively).

Similarly, higher levels of virus-specific antibodies were induced by each of the B2 immunogens when these were inoculated in the presence of adjuvant (virus p = 0.0003; wtVLP p = 0.207; rsVLP p<0.0001), but this was not the case for the C4 (18/116) immunisation group (p = 0.6703). Both genogroup B2 virus and rsVLP immunisation generated antibodies that detected B2 virus in ELISA, and performed better than the genogroup C4 immunogen (18/116) (p = 0.0015 & p = 0.0002, respectively). This improvement over C4 (18/116) immunisation was not as pronounced for the wtVLP immunogen (p = 0.1194) (Fig 8A). In each instance, adjuvanted genogroup B2 immunisation, whether with virus, wtVLP or rsVLP, generated higher neutralising antibody titres than the 18/116 genogroup C4 control (p = 0.0215, p = 0.0092, p = 0.0022, respectively) (Fig 8b). The presence of adjuvant, particularly with the VLP immunisations, was a critical factor in the generation of both reactive antibodies and neutralising antibodies against genotype B2 virus.

## Discussion

The functional importance of the antigenic conformation of PV vaccines is well understood; in order to generate long-lived protective immunity against PV it appears to be necessary to vaccinate with NAg particles. It is for this reason that the antigenic metric for quality control of PV vaccines is the NAg (DAg) unit. All PV vaccines contain minor amounts of HAg particles, which are irrelevant for the generation of protective immunity. However, several groups have described the generation of neutralising antibodies against EVA71 VLPs which have been derived from a wt sequence, presumably in the HAg conformation [18–20]. To directly address whether there is a significant immunological difference between NAg and HAg vaccines against EVA71 in a murine model we generated wtVLPs and rsVLPs and assessed particle stability, determined the structure of the stabilised particles and carried out comparative immunisation studies.

Consistent with previous reports, we found that EVA71 VLPs can be produced efficiently in *P. pastoris* (Fig S1). Both the wt and rsVLP constructs produced particles that could be purified following established protocols [30]. The wtVLPs were predominantly HAg. In contrast, the rsVLP population was primarily NAg (87%) and stable to >44°C (Fig 2, Fig 3, Table S1).

High-resolution structural data generated from the rsVLPs provide important insight into a potential mechanism for particle stabilisation. Particles were separated into two classes, with the major class containing approximately 90% of the particles and resolved to 2.4 Å, and the minor population resolved to 3.4 Å. The major class displayed features characteristic of the NAg conformation; density for a sphingosine-like pocket factor, a narrow opening at the 2-fold symmetry axis, organised internal density for much of the VP1 N-terminal region, and a VP3 GH-loop that extends toward the 2-fold axis at the quasi-3-fold axis (Fig 4A, Fig 4B, Fig 5).

Curiously, the minor particle class was not entirely consistent with either the NAg or HAg forms of EVA71 and showed weaker localised density in the inter-pentamer regions of the capsid (Fig 4C, Fig 4D). While the models for the major and minor populations were remarkably similar (rmsd 0.798), the primary notable difference was the ability to resolve an additional 14 residues within the N-terminal region of VP1 (aa 58-70). Despite the differences between the major and minor rsVLPs at the inter-pentamer interfaces, the diameter of the particles (approximately 30.6 nm), the presence of sphingosine-like density within the pocket (Fig S5), the conformation of the VP1 β-barrel and the orientation of the phenyl group of VP1 F233 all suggest retention of the NAg conformation and a lack of particle expansion for both major and minor rsVLP classes [22].

The precise role for each individual mutation in particle stabilisation is not clear, although we believe it is likely that the mutations are functioning cooperatively to provide the stabilisation phenotype. The position of the VP3 C-terminal chain mutation I235M atop the canyon at the base of the VP1 β-barrel is in close proximity to VP1 Y116, although this does not appear to interact directly with Y116 (Fig 5). However, VP1 Y116 is in a part of the BC-region that forms a small helix at the top of the canyon, and rotation of this sidechain between the NAg and HAg forms may contribute to separation of the B and C strands within the VP1 β-barrel and help to facilitate release of the stabilising pocket factor in HAg particles. The presence of a cysteine in place of this tyrosine (Y116C) may reduce the tendency for β-barrel collapse and contribute to maintenance of the pocket in a ‘closed’ state.

In the presence of the K162I mutation, the inability to form salt bridge between VP1 D110 and K162 (as observed in wt virions) results in several minor changes to the opening of the pocket and ultimately a ~15% reduction in the diameter of the opening in the direction perpendicular to the canyon. These minor changes include movement of the carboxylate group of D110 such that it sits further over the mouth of the pocket atop the canyon in the rsVLPs. A shift in position of the Cα of D110-I111-D112 (0.653 Å rmsd) residues also results in slight narrowing of the width of the opening parallel with the canyon (~0.45 Å)(Fig 6). Together, these minor changes result in a significant narrowing of the mouth of the pocket that could hinder the release of the lipid “pocket factor” and thus contribute to increased particle stability (Fig 6).

The VP1 K162I mutation may also stabilise the rsVLP indirectly by destabilizing the HAg form. VP1 K162 forms a salt-bridge with VP1 D110 in the NAg particles, helping to stabilise the position of the C-terminal end of the VP1 B-strand and maintain the stabilising pocket factor within the assembled particle. Upon particle expansion, VP1 K162 is no longer able to form this salt bridge, instead this loop extends away from the collapsed pocket toward the VP3 GH-loop (PDB-3VBS and PDB-3VBU structures [22]). Additionally, this GH-loop in the expanded form is extended away from the shell of the capsid and an Asp within this loop may form a hydrated salt bridge with VP1 K162 from an adjacent protomer. In the presence of the K162I mutation, this salt bridge cannot form, potentially leading to a destabilisation of HAg, favouring retention of the NAg conformation (Fig 5). Notably, the VP3 GH-loop was retained in the NAg conformation for the minor class of rsVLP, suggesting that these particles may be intermediates between NAg and HAg (stabilised by rsVLP mutations). However, it remains possible that they represent an alternative off-pathway conformation e.g. NAg particles with assembly defects.

To correlate the structural and antigenic differences described above with immunogenicity, we carried out comparative immunisation studies. Unsurprisingly, in the presence of adjuvant, sera from each of the B2 genotype experimental immunisation groups performed better than their respective unadjuvanted counterparts, inducing antibodies reactive with genogroup B2 virus (Fig 8A). Interestingly, in each case immune sera were better at detecting rsVLPs than wtVLPs. This could suggest that that rsVLPs are more antigenic, despite showing no increase in immunogenicity compared to the wt control immunogens. Alternatively, the CT11F9 antibody used to quantify VLPs may underestimate the total amount of NAg particles present in the rsVLP sample (Fig 7 and Fig S6).

Consistent with previous observations describing the generation of anti-EVA71 antibodies in mice, we have shown that immunisation with predominantly NAg or HAg particles generated neutralising antibody titres similar to those induced by a wt B2 virus immunogen (Fig 8). Curiously, in the presence of adjuvant, only a minor improvement in neutralising antibody titre was detected for the B2 virus and the C4 18/116 immunisation groups (p = 0.0538 & p = 0.4634, respectively), suggesting that the RNA genomes contained within the particles may be immunostimulatory. Genotype-specific differences were demonstrated, as each of the adjuvanted B2 immunogens generated higher reactive and neutralising titres against B2 virus compared to the genogroup C4 18/116 immunogen (Fig 7, Fig 8).

Collectively, these data indicate that (under the conditions tested here and in a murine model of anti-EVA71 immunisation), that there was no significant advantage afforded to either NAg or HAg immunogens within a given genogroup. A slight improvement in anti-virus endpoint titre was observed for rsVLPs compared to wtVLPs (p = 0.0621) in the presence of adjuvant, although this did not correlate with an improvement in neutralising antibody titres (Fig 7 & Fig 8). When adjuvanted, VLPs and virions have equivalent immunogenicity, however it is likely that there is a genotype-specific preference in the generation of total antibody titres and neutralising antibody titres. We note that, given the characteristic differences between mouse and human antibodies, differences in the efficacy of NAg or HAg immunogens may be more apparent in alternative animal models. Additionally, the minor differences we observe in responses induced by immunisation warrant further investigation, particularly for a more complete characterisation of protective responses which do not rely upon direct virus neutralisation (e.g. ADCC, complement activation etc.), and a characterisation of the performance of these vaccines in a challenge model, where additional host factors can contribute to anti-viral responses.

Importantly, the data generated in this study improves our understanding of the mechanism of antigenic conversion in EVA71. In immunisation experiments wt and rsVLPs behaved similarly to virus. However, they are cheaper and safer to manufacture and EVA71 VLPs may therefore offer significant advantages for vaccine manufacturing in the future.

## References

## Conflict of interest

The Authors declare no conflict of interest.

## Acknowledgements

We would like to thank Dr Tong Cheng (Xiamen University) for sharing the CT11F9 antibody.

## Funding

We gratefully acknowledge support from The UK Medical Research Council MR/P022626/1 (NJK, NJS, DJR, AJM, JM and AM) and support from the NIH R01 AI 169457-0 (NJK, NJS, DRJ). NJK and KG hold fellowships from the Wellcome Trust ISSF (204825/Z/16/Z) and JSS holds a Wellcome Trust studentship (102174/B/13/Z). JMH held a Leverhulme Trust Visiting Professor Fellowship at the University of Leeds.

## Author contributions

NJK, NJS, DJR and AJM conceived and planned experiments. Funding was sourced by NJS, DJR, AJM, JM, NJK, JMH. NJK generated VLPs, using mutations selected by MS. NJK and KG antigenically characterised VLPs. NJK and JSS generated structures of stabilised VLPs and performed structural analysis of stabilised VLPs. AM, AT, EP and HF carried out immunisations and performed neutralisation assays. NJK assessed sera for reactive titres. NJK prepared the initial manuscript, and all authors were involved in review of the data and review of the manuscript.

## Data availability

The atomic model for the recombinant stabilised VLP is available through PDB ID 8C6D and the density map is available with EMDB ID 16450.

## Supplementary material

**Figure S1:**
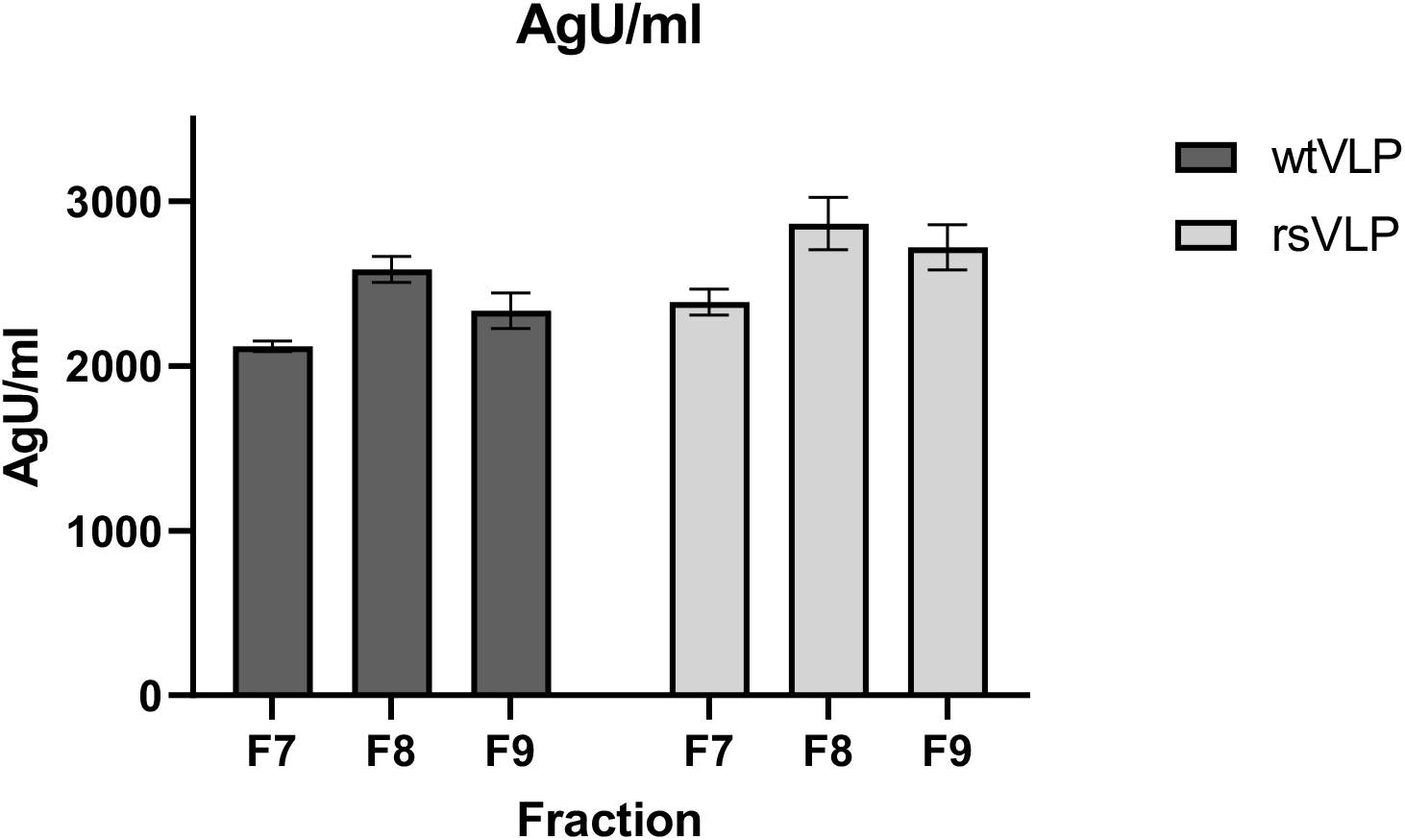
Antigen concentration (AgU/ml) of peak VLP fractions determined by the CT11F9 antibody relative to the 18/116 antigen standard.

**Figure S2:**
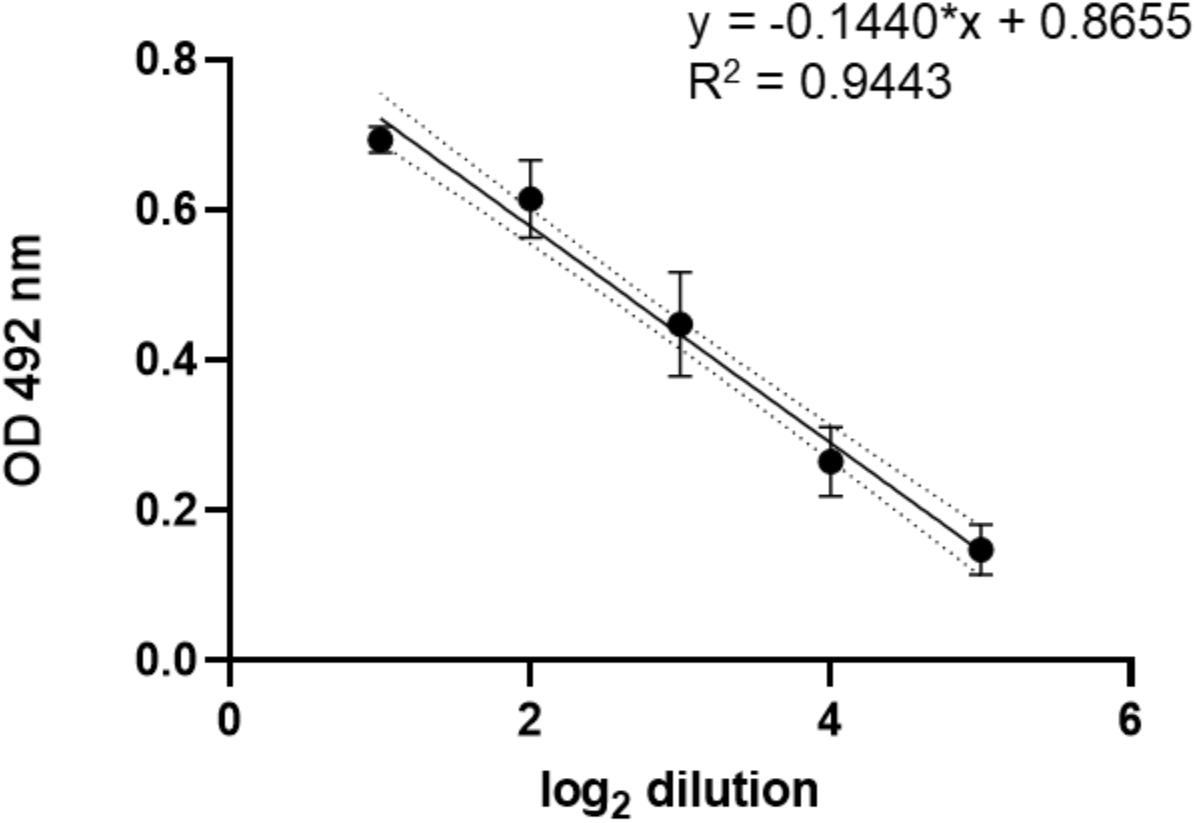
Relationship between OD and log_2_ dilution of antigen, used to calculate the proportion of HAg in samples.

**Figure S3:**
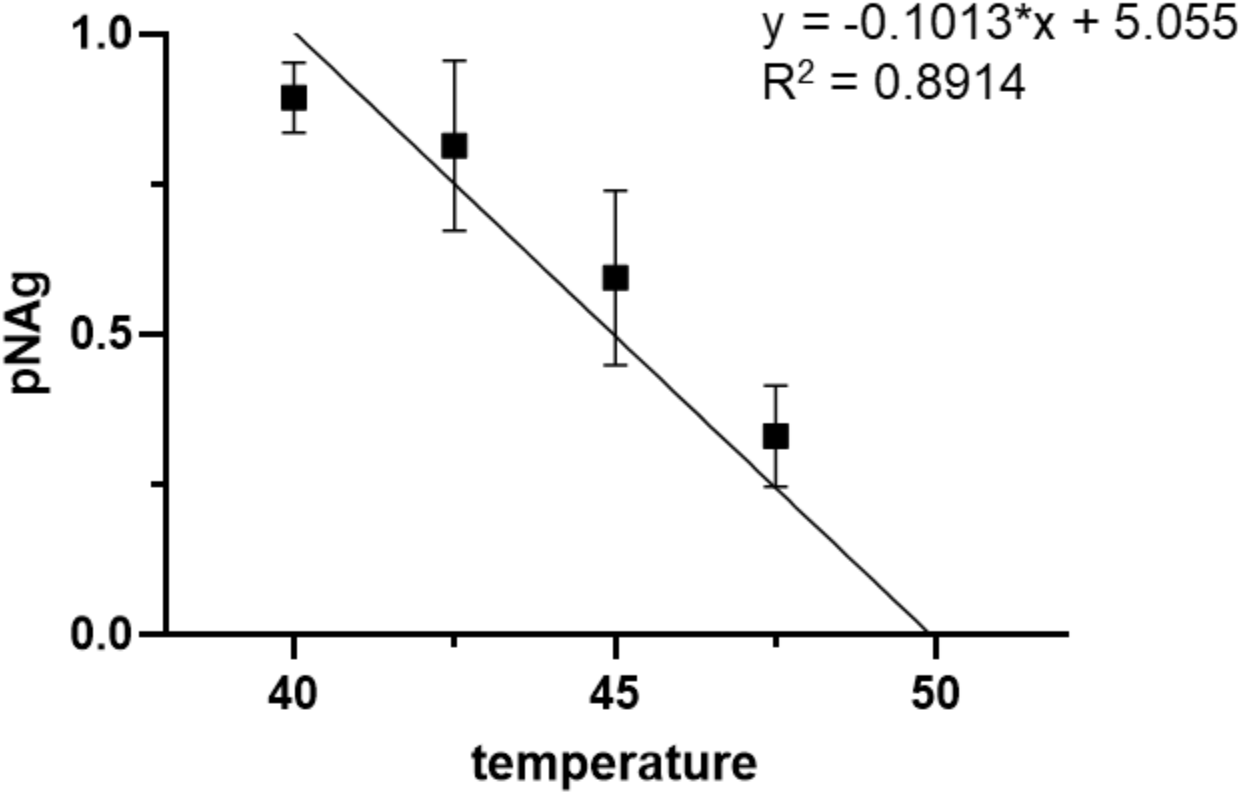
Linear regression to estimate antigenic conversion temperature of rsVLPs between 40°C and 50°C.

**Figure S4:**
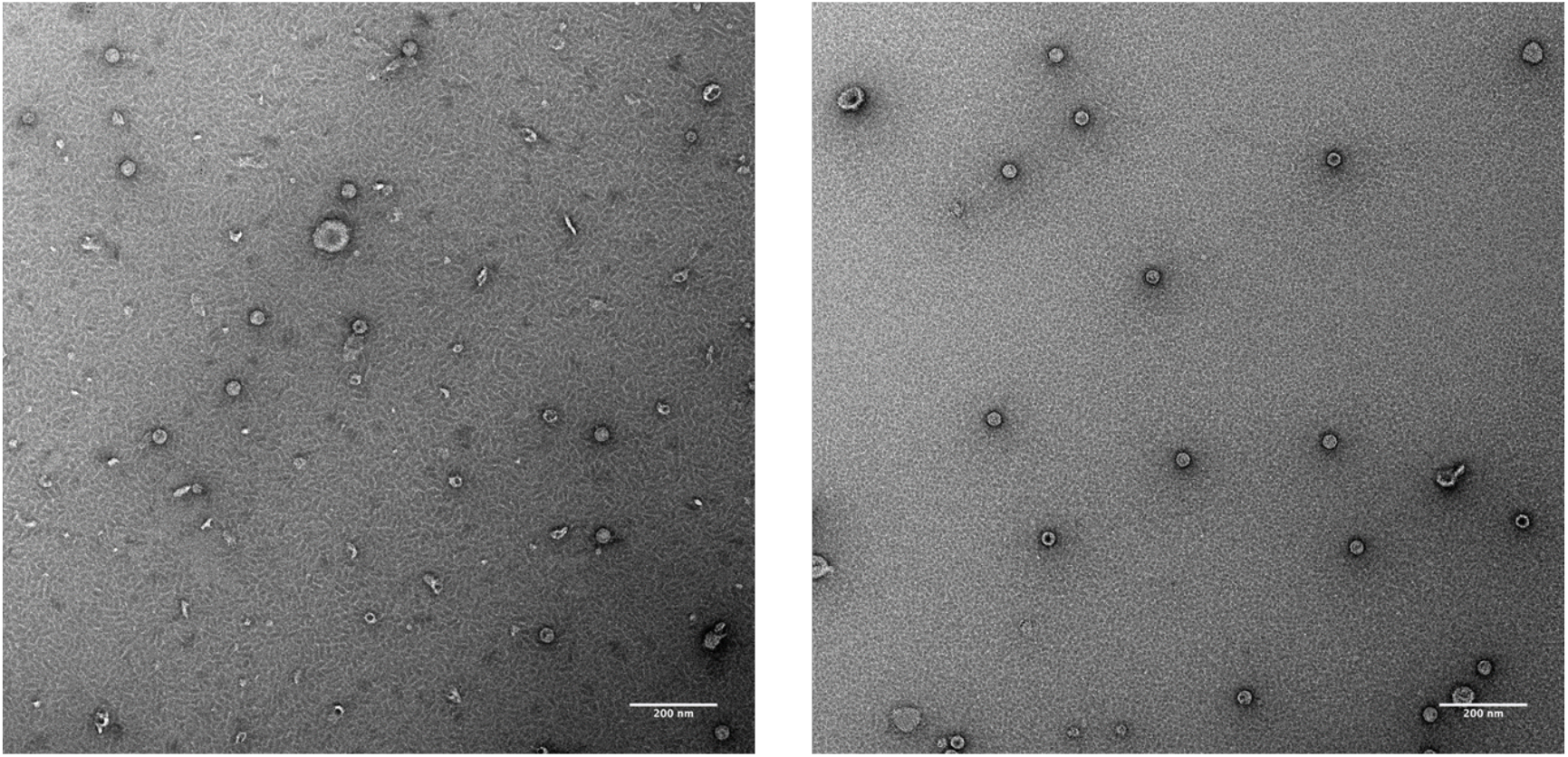
Negative stain TEM (using 2% UA) of samples of wtVLPs (left) and rsVLPs (right), scale bar = 200 nm.

**Figure S5:**
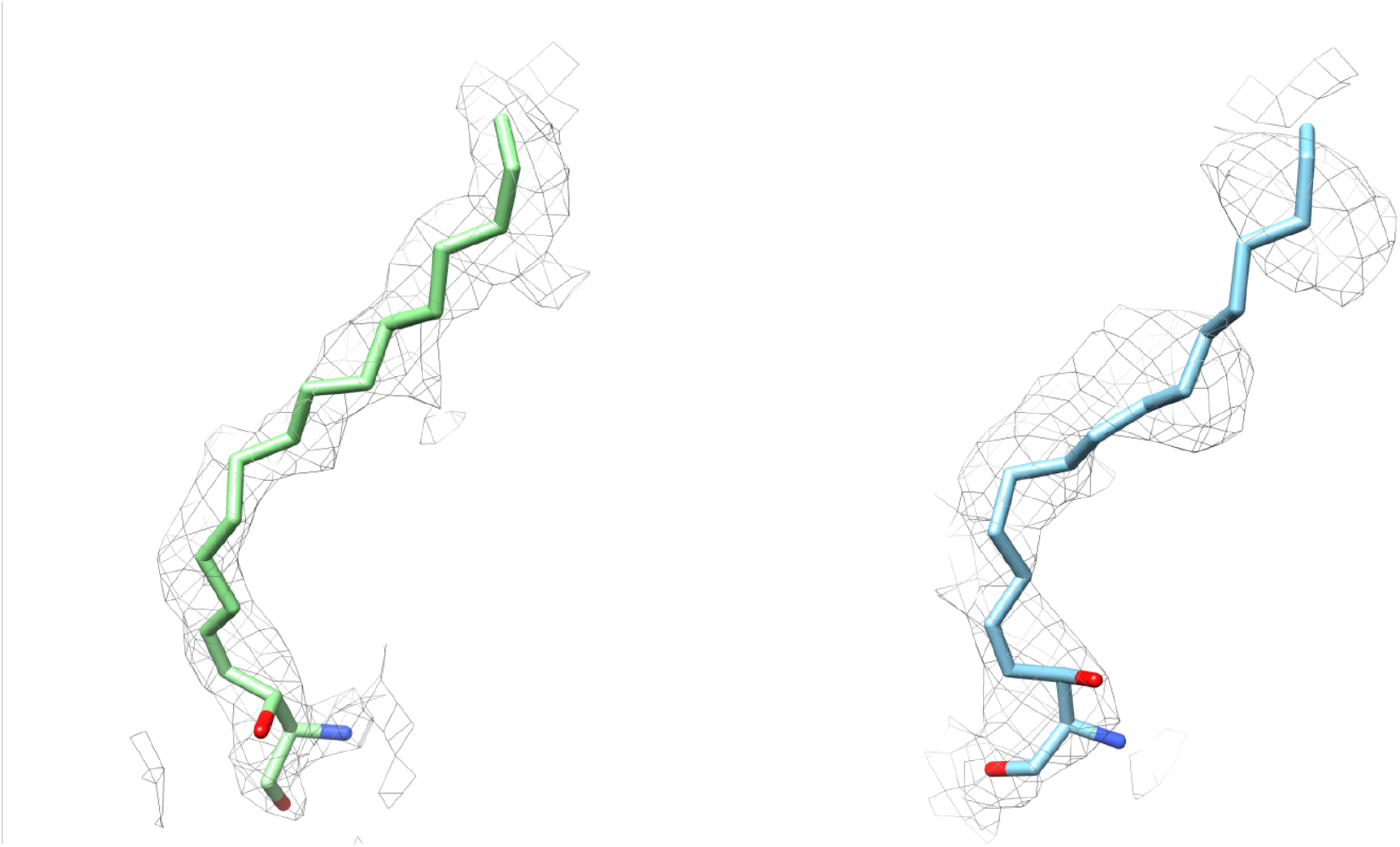
Density indicating occupancy of the pocket, present in the major population (left) and the minor population (right) shown at 2.5 rmsd.

**Figure S6:**
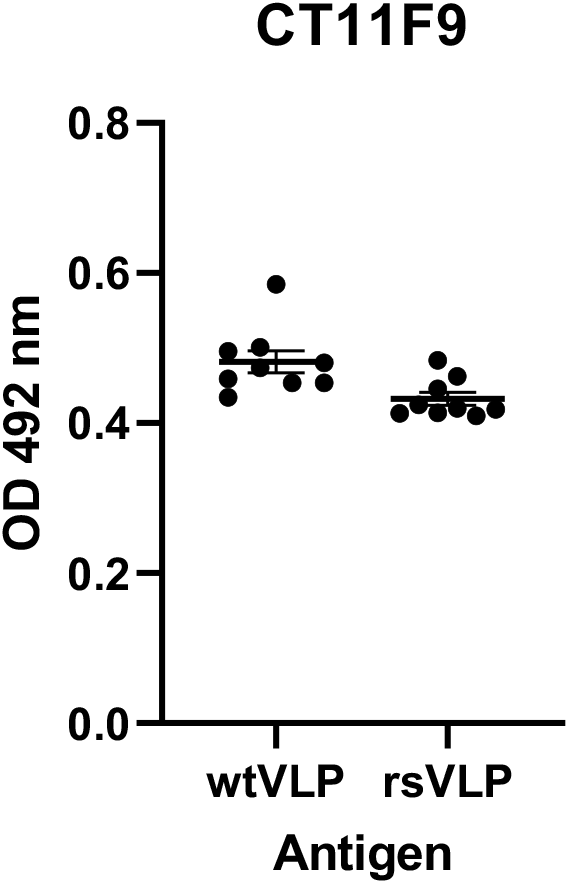
Reactivity of VLP (captured on ELISA plates) with CT11F9 antibody.

**Table S1:** Antigen composition of samples at a range of temperatures

$$\text{proportion HAg} = 2^{(OD\ 4^{\circ}C - OD\ 55^{\circ}C)/\text{incline}}$$
| Temperature (°C) | pHAg rsVLP | pNAg rsVLP | pHAg wtVLP | pNAg wtVLP |
| --- | --- | --- | --- | --- |
| 4 | 0.13 | 0.87 | 1.11 | -0.11 |
| 30 | 0.11 | 0.89 | 0.80 | 0.20 |
| 35 | 0.10 | 0.90 | 0.69 | 0.31 |
| 37.5 | 0.11 | 0.89 | 0.84 | 0.16 |
| 40 | 0.14 | 0.86 | 0.97 | 0.03 |
| 42.5 | 0.31 | 0.69 | 1.16 | -0.16 |
| 45 | 0.46 | 0.54 | 1.18 | -0.18 |
| 47.5 | 0.86 | 0.14 | 1.32 | -0.32 |
| 50 | 1.10 | -0.10 | 1.54 | -0.54 |
| 52.5 | 1.07 | -0.07 | 1.17 | -0.17 |
| 55 | 1.00 | 0.00 | 1.00 | 0.00 |
| 60 | 0.80 | 0.20 | 0.73 | 0.27 |

**Table S2:**

|  | rsVLP |
| --- | --- |
| Microscope | FEI Titan Krios |
| Detector mode | Counting |
| Camera | Falcon IV |
| Voltage (kV) | 300 |
| Pixel size (Å) | 0.91 |
| Nominal magnification | 130,000× |
| Exposure time (s) | 5 |
| Total dose (e<sup>-</sup>/Å<sup>2</sup>) | 31 |
| Number of fractions | 42 |
| Defocus range (μm) | -0.8 to -2.9 |
| Number of micrographs | 20,419 |
| Acquisition software | Thermo Scientific EPU |

**Table S3:**
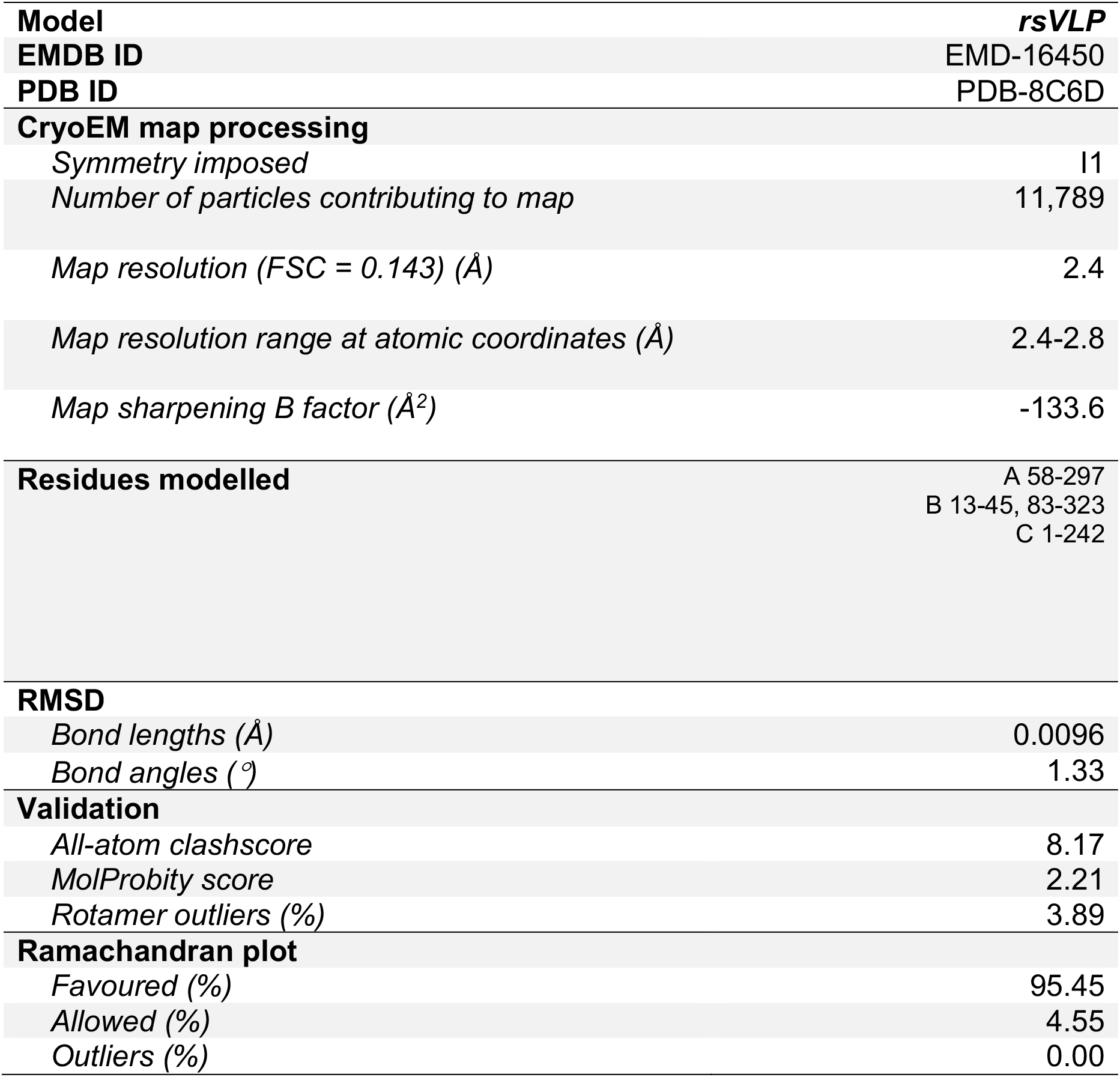

## Notes

### Competing Interest Statement

The authors have declared no competing interest.

